# Enhancer control of miR-155 expression in Epstein-Barr virus infected B cells

**DOI:** 10.1101/311886

**Authors:** C. David Wood, Thomas Carvell, Andrea Gunnell, Opeoluwa O. Ojeniyi, Cameron Osborne, Michelle J. West

**Affiliations:** School of Life Sciences, University of Sussex, Falmer, Brighton, UK; Department of Medical and Molecular Genetics, King’s College London School of Medicine, Guy’s Hospital, London, UK

## Abstract

The oncogenic microRNA-155 (miR-155) is the most frequently upregulated miRNA in Epstein-Barr virus (EBV)-positive B cell malignancies and is upregulated in other non-viral lymphomas. Both the EBV nuclear antigen 2 (EBNA2), and B cell transcription factor, interferon regulatory factor 4 (IRF4) are known to activate transcription of the host cell gene from which miR-155 is processed (*miR-155HG,* BIC). EBNA2 also activates *IRF4* transcription indicating that EBV may upregulate miR-155 through direct and indirect mechanisms. The mechanism of transcriptional regulation of *IRF4* and *miR-155HG* by EBNA2 however has not been defined. We demonstrate that EBNA2 can activate *IRF4* and *miR-155HG* expression through specific upstream enhancers that are dependent on the Notch signaling transcription factor RBPJ, a known binding partner of EBNA2. We demonstrate that in addition to activation of the *miR-155HG* promoter, IRF4 can also activate *miR-155HG* via the upstream enhancer also targeted by EBNA2. Gene editing to remove the EBNA2- and IRF4-responsive *miR-155HG* enhancer located 60 kb upstream of *miR-155HG* led to reduced *miR155HG* expression in EBV-infected cells. Our data therefore demonstrate that specific RBPJ-dependent enhancers regulate the IRF4-miR-155 expression network and play a key role in the maintenance of miR-155 expression in EBV-infected B cells. These findings provide important insights that will improve our understanding of miR-155 control in B cell malignancies.

**IMPORTANCE:** MicroRNA-155 (miR-155) is expressed at high level in many human cancers particularly lymphomas. Epstein-Barr virus (EBV) infects human B cells and drives the development of numerous lymphomas. Two EBV-encoded genes (LMP1 and EBNA2) upregulate miR-155 expression and miR-155 expression is required for the growth of EBV-infected B cells. We show that the EBV transcription factor EBNA2 upregulates miR-155 expression by activating an enhancer upstream from the miR-155 host gene (*miR-155HG*) from which miR-155 is derived. We show that EBNA2 also indirectly activates *miR-155* expression through enhancer-mediated activation of *IRF4.* IRF4 then activates both the *miR-155HG* promoter and the upstream enhancer, independently of EBNA2. Gene editing to remove the *miR-155HG* enhancer leads to a reduction in *miR-155HG* expression. We therefore identify enhancer-mediated activation of *miR-155HG* as a critical step in promoting B cell growth and a likely driver of lymphoma development.

## INTRODUCTION

MicroRNAs (miRNAs) are a class of highly conserved, non-coding RNA molecules of 18-25 nucleotides in length that play an important role in post-transcriptional gene control. MiRNAs hybridize to target mRNAs, often in the 3’ untranslated region, and promote their degradation and/or inhibit their translation. MiRNAs can be transcribed from specific promoters or processed from coding or non-coding gene transcripts. Deregulation of miRNA expression is implicated in the pathogenesis of many diseases, including a diverse range of human cancers and the term oncomiR is used to describe miRNAs with tumor-promoting properties (1).

The miR-155 oncomiR was originally discovered as a non-coding RNA within the B cell integration cluster (*BIC*) gene (2). *Bic* was previously identified as a proto-oncogene activated by proviral insertion in avian leucosis virus-induced lymphomas (3, 4). The miR-155 locus is highly conserved across species and in humans lies within the third exon of *BIC* (miR-155 host gene; *miR-155HG).* MiR-155 appears to play a key role in the regulation of B lymphocyte function. Transcription of *miR-155HG* is activated upon B cell receptor signaling and in murine models dysfunction or loss of miR-155 in B lymphocytes causes a severe decrease in antibody-induced signaling (5, 6). Overexpression of miR-155 in mice results in the development of precursor B lymphoproliferative disorders and B cell lymphomas (7). MiR-155 expression is highly upregulated in a number of human lymphomas including Hodgkin’s and diffuse large cell B-cell lymphoma (5, 8, 9). The basis of the oncogenic activity of miR-155 has not been fully elucidated however a number of target genes that regulate B cell proliferation and survival have been identified. These include transcription regulators, receptors and signaling pathway components e.g. *HDAC4, PIK3R1, SMAD5, SHIP1, PU.1, BCL2* and *C/EBPβ* (10, 11).

Epstein-Barr virus (EBV) immortalizes human B lymphocytes and is associated with the development of numerous lymphomas including Burkitt’s, Hodgkin’s and diffuse large B cell (DLBCL). MiR-155 expression is upregulated on B cell infection by EBV (12). In *in vitro* EBV transformed B cell lines (lymphoblastoid cell lines; LCLs) and an EBV-positive DLBCL cell line, loss of miR-155 expression inhibits cell growth and induces apoptosis indicating that miR-155 expression is important for transformed B cell survival (13). MiR-155 expression in LCLs appears to attenuate high levels of NF-κB signaling and this may help promote B cell proliferation and prevent apoptosis (14). Consistent with a key role for gene regulation by miR-155 in viral-induced oncogenesis, the oncogenic herpesviruses Kaposi’s sarcoma herpesvirus and Marek’s disease herpes virus encode miR-155 mimics in their viral genomes (15-17).

Two EBV genes essential for B cell transformation upregulate miR-155 expression; the constitutively active CD40 receptor mimic, latent membrane protein 1 (LMP1) and the viral transcription factor, Epstein-Barr virus nuclear antigen 2 (EBNA2) (13, 14). Expression of either LMP1 or EBNA2 independently activates transcription of *miR-155HG* (14). Upregulation of AP-1 and NF-κB activity by LMP1 appears to play an important role in activation of the miR-155 promoter in EBV-infected cells (18, 19). The mechanism of EBNA2 activation of miR-155 has not been demonstrated. EBNA2 is required for B cell immortalization by EBV and activates all viral gene promoters, including LMP1, so indirect activation of miR-155 via upregulation of LMP1 is a likely consequence of EBNA2 expression (20, 21). However, EBNA2 also deregulates host gene transcription by binding to promoter and enhancer elements (22, 23). Enhancer and super-enhancer activation by EBNA2 appears to be widespread in the B cell genome (23-25). For example, EBNA2 activation of the *MYC* proto-oncogene is directed by the targeting of upstream enhancers and modulation of enhancer-promoter looping (22, 26). EBNA2 does not bind DNA directly and associates with viral and cellular gene regulatory elements through its interactions with cellular transcription factors that include RBPJ, PU.1 and EBF1 (27).

An EBNA2-bound super-enhancer postulated to control miR-155 expression was identified in LCLs based on the binding of a number of EBV transcription factors (EBNA2, EBNA3A, EBNA3C and EBNA-LP), binding of NF-κB subunits and broad and high histone H3 lysine 27 acetylation signals (25). However, the original region identified actually comprises the highly expressed 20 kb *miR-155HG* transcription unit from which miR-155 is derived. A subsequent study using RNA polymerase II (RNA pol II) chromatin interaction analysis by paired-end tag sequencing (ChIA-PET) found that RNA pol II associated with a number of EBNA2-bound promoter, enhancer and super-enhancer regions upstream of *mIR-155HG* that formed links with the *mIR-155HG* promoter (28). Whether EBNA2 can activate *miR-155HG* transcription via the *miR-155HG* promoter or these putative enhancer elements however has not been investigated.

MiR-155 expression is also activated by interferon regulatory factor 4 (IRF4) through an interferon-stimulated response element (ISRE) in the *miR-155HG* promoter (29).

Interestingly, IRF4 levels are highly upregulated in EBV infected cells and like miR-155, *IRF4* is also induced by both LMP1 and EBNA2 (30). As a result, *IRF4* and miR-155 levels correlate in EBV infected cells. In addition to the potential indirect effects of EBNA2 on *IRF4* expression via LMP1 upregulation, conditional expression of EBNA2 in the presence of protein synthesis inhibitors also demonstrates that *IRF4* is a direct target gene of EBNA2 (31). The mechanism of EBNA2 activation of *IRF4* has not been demonstrated. IRF4 expression is essential for the growth and survival of LCLs and apoptosis induced by IRF4 depletion can be partially rescued by expression of miR-155 (29, 32). This indicates that the upregulation of miR-155 by IRF4 may be a key component of its essential role in promoting LCL growth.

To obtain important information on how the IRF4/miR-155 expression network is controlled by EBV, we investigated the role of putative upstream EBNA2-bound enhancer elements in the regulation of *miR-155HG* and *IRF4* expression. At both gene loci we identified specific EBNA2-bound enhancer elements that activate transcription of their respective promoters in an RBPJ-dependent manner. Deletion of the EBNA2-responsive *miR-155HG* enhancer resulted in a decrease in *mIR-155HG* transcription in EBV-infected cells demonstrating its importance for the maintenance of miR-155 expression. These data identify key enhancer elements utilized by EBV for the control of two genes critical for B cell growth that is relevant to the study of miR-155 and *IRF4* deregulation in other tumor contexts.

## RESULTS

### A miR-155HG upstream enhancer is activated by EBNA2 through RBPJκ

To obtain information on regulatory elements that may control miR-155 expression, we examined *miR-155HG* promoter interaction data obtained by the genome-wide chromosome conformation technique, capture Hi-C (CHi-C) (Figure 1). In both the GM12878 LCL and in CD34+ hematopoietic progenitor cells, the *miR-155HG* promoter interacts with three main upstream regions marked by high levels of H3K27ac indicating transcription regulatory function. These include two intergenic regions and an intragenic region proximal to the promoter of the *LINC00158* non-coding RNA gene (Figure 1). The same *miR-155HG* interacting regions were also detected by RNA pol II ChIA-PET (28). Interestingly, CHi-C data demonstrates that the miR-155 genomic locus within exon 3 of *miR-155HG* interacts at a much lower frequency with the two intergenic regions (Figure 1). This suggests that these interactions more frequently involve the *miR-155HG* promoter, consistent with a role in regulating transcription. The miR-155 genomic locus does however interact with the *LINC00158* promoter proximal region, consistent with a gene to gene looping interaction between *miR-155HG* and *LINC00158* (Figure 1). The *miR-155HG-LINC00158* interaction is also the main interaction detected in this region by ChiA-PET for the chromatin organizing factor CTCF, suggesting it may be involved in domain organization rather than *miR-155HG* promoter regulation (28). Our EBNA2 chromatin immunoprecipitation ChIP-sequencing data from the same LCL used for CHi-C detects the highest EBNA2 binding at two sites within the most proximal intergenic region (24) (Figure 1). We therefore investigated the role of these two EBNA2-bound putative enhancers in the regulation of *miR155-HG.*

**FIG 1.**
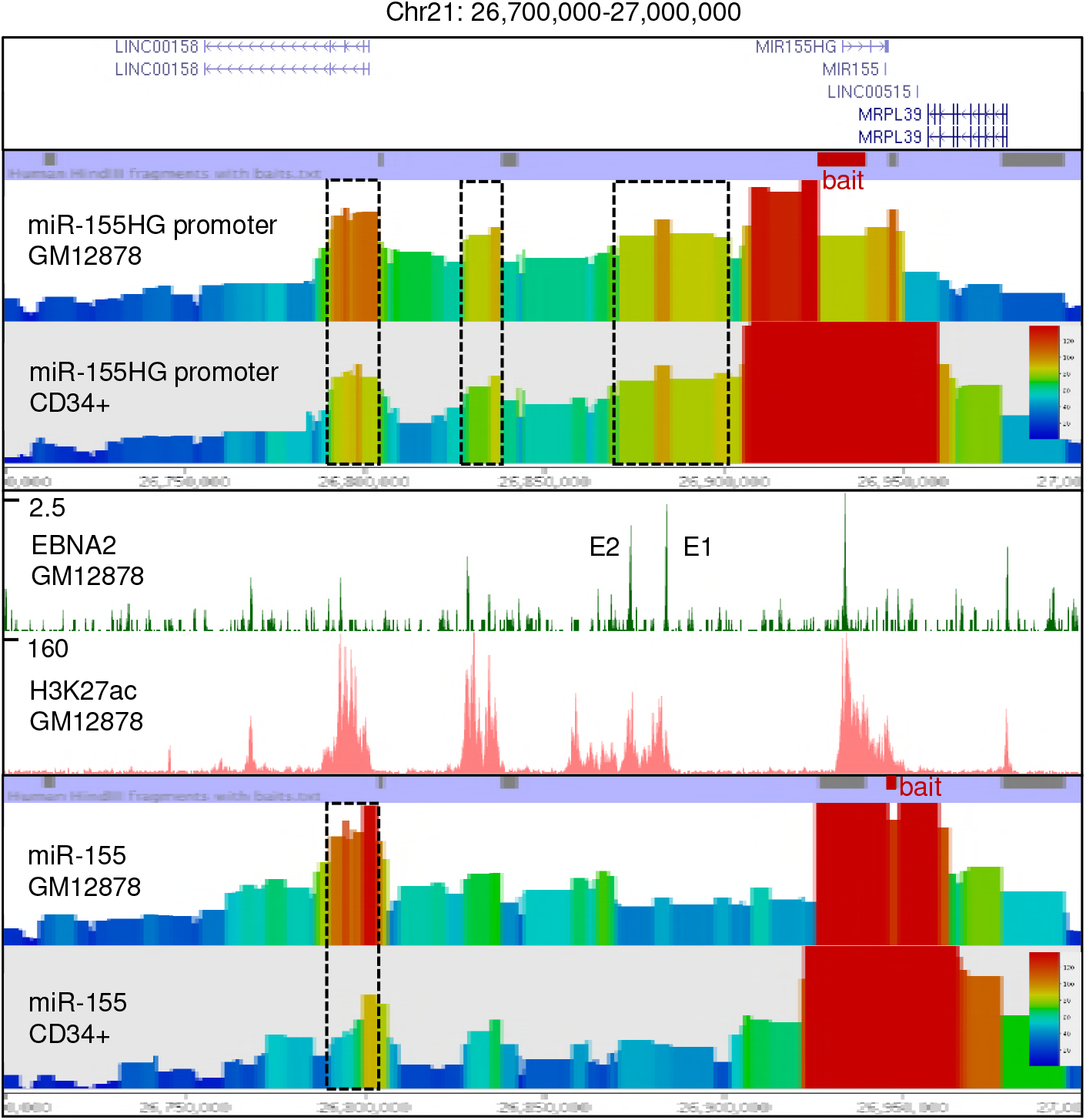
Chromosome interactions and EBNA2 binding at the *miR-155HG* locus on chromosome 21. Capture Hi-C interaction data obtained using *HindIII* fragments encompassing the *miR-155HG* promoter (top panel) or the miR-155 genomic locus (bottom panel) as bait (red boxes above CHi-C data bar charts). Bar charts show the geometric mean of sequencing reads rainbow colored by read frequency according to the scale bar. Interacting fragments were captured from Hi-C libraries generated from the EBV infected LCL GM12878 or CD34+ progenitor cells (50). The main interacting regions are shown in boxes with dashed lines. EBNA 2 ChIP-sequencing reads in GM12878 cells (24) and H3K27Ac signals in GM12878 from ENCODE are shown (middle panel). The positions of the two main EBNA2-bound putative enhancer regions are indicated (E1 and E2).

We generated luciferase reporter plasmids containing the *mIR-155HG* promoter and one or both of the enhancer elements (E1 and E2). Reporter assays carried out in the EBV negative B cell line DG75 in the absence or presence of transient EBNA2 expression demonstrated that EBNA2 had no effect on the *miR-155HG* promoter but activated transcription up to 7.3-fold when a region encompassing both E1 and E2 was inserted upstream of the promoter (Figure 2A). The level of activation was similar to that observed for the EBNA2 responsive EBV C promoter (Figure 2B). When testing each enhancer separately, we found that the presence of E1 alone did not convey EBNA2 responsiveness, but it increased basal transcription levels compared to the promoter alone by approximately 2-fold (Figure 2A). This indicates that E1 has EBNA2-independent enhancer function. EBNA2 activated transcription via E2 alone up to 10.8-fold indicating that E2 is an EBNA2-responsive enhancer (Figure 2A). Interestingly, the presence of E2 decreased basal transcription levels approximately 5-fold compared to the promoter alone (Figure 2A). This is consistent with the presence of repressive elements in the enhancer that can limit basal transcription activity, a feature we observed previously for the main EBNA2-responsive enhancer at *RUNX3* (24). As a result, the overall level of transcription in the presence of E2 was lower than that in the presence of E1 and E2 combined (Figure 2A). Since EBNA2 upregulates *IRF4* and IRF4 is a known activator of *miR-155HG,* we investigated whether the effects of EBNA2 in these reporter assays may be indirect and the result of increased endogenous IRF4 expression. We found that transient expression of EBNA2 did not increase endogenous IRF4 expression (Figure 2A). We conclude that EBNA2 independent and EBNA2-dependent enhancers regulate *miR-155HG* transcription in EBV-infected cells and that EBNA2 activates transcription directly via association with a specific *miR-155HG* enhancer.

**FIG 2.**
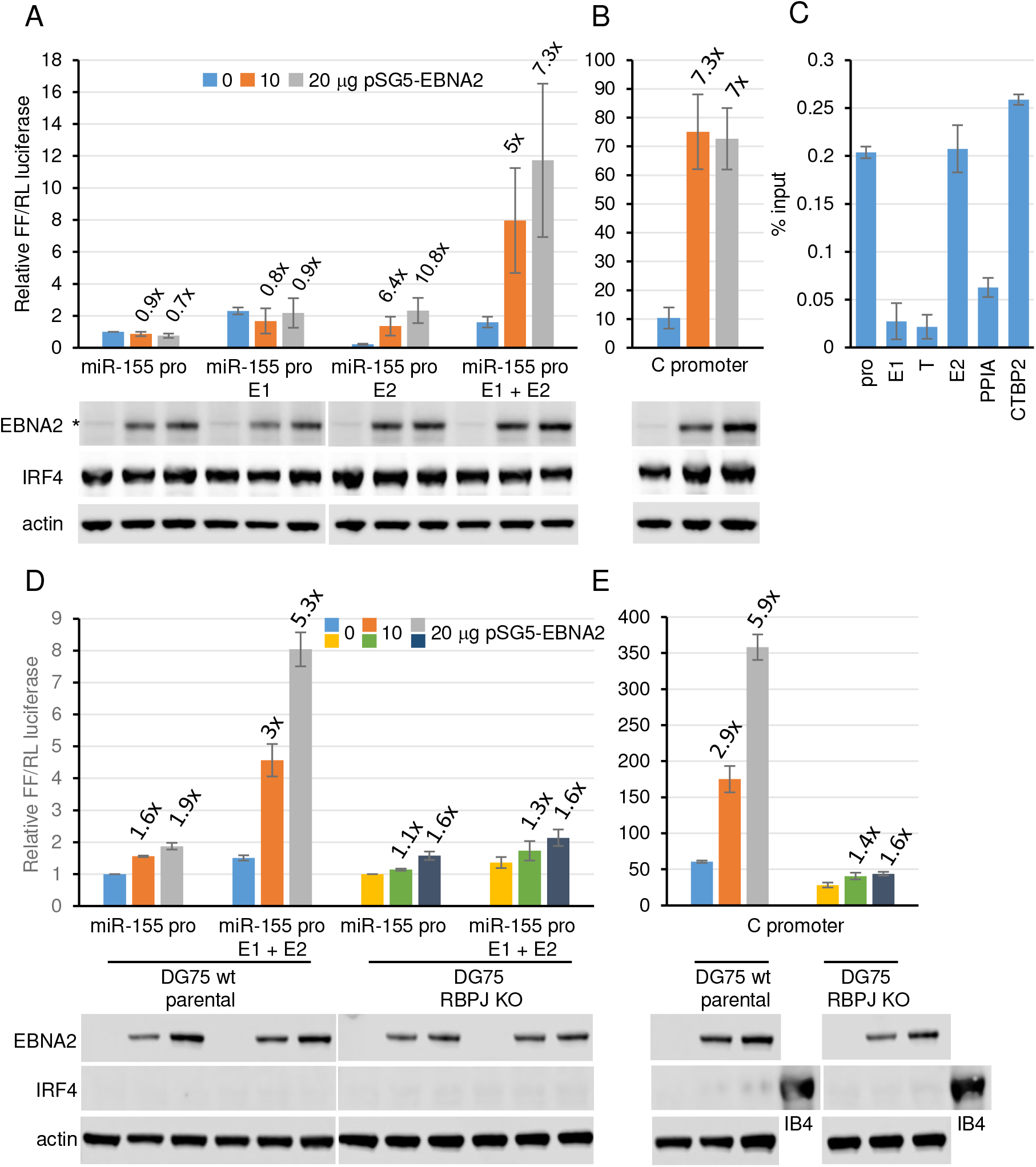
The effects of EBNA2 on *miR-155HG* promoter and enhancer elements. (A) *mIR-155HG* luciferase reporter assays in the presence or absence of EBNA2. DG75 cells were transfected with 2 μg of pGL3 firefly luciferase reporter constructs containing the *miR-155HG* promoter either alone or in the presence of enhancer E1, E2 or both E1 and E2. Assays were carried out in the absence or presence of 10 or 20 μg of the EBNA2-expressing plasmid pSG5-EBNA2 and 0.5 μg of Renilla luciferase control plasmid (pRL-TK). Firefly luciferase signals were normalized to Renilla luciferase signals and expressed relative to the signal obtained for the *miR-155HG* promoter in the absence of EBNA2. Results show the mean of three independent experiments plus or minus standard deviation. Fold activation by EBNA2 relative to the signal obtained for each construct in the absence of EBNA2 is shown above each bar. Western blot analysis of EBNA2 and IRF4 expression is shown below each bar chart, with actin providing a loading control. All blots shown were probed at the same time with the same batch of antibody solution and for each protein show the same exposure. They are therefore directly comparable, but have been cut and placed to align with the respective luciferase assay graphs. The asterisk shows the position of a non-specific band visible on longer exposures of EBNA2 blots. (B) EBNA2 activation of an EBV C promoter reporter construct was used as a positive control. (C) ChIP-QPCR analysis of RBPJκ binding at the *miR-155HG* locus in GM12878 cells. Precipitated DNA was analysed using primer sets located at the promoter, E1, E2 and in a trough between E1 and E2 (T). EBNA2 binding at the transcription start site of *PPIA* and at the previously characterised *CTBP2* binding site were used as negative and positive binding controls, respectively. Mean percentage input signals, after subtraction of no antibody controls, are shown plus or minus standard deviation for three independent ChIP experiments. (D) Luciferase reporter assays carried out using the *miR-155HG* promoter or *miR-155HG* E1 and E2 construct in DG75 wt parental cells that lack IRF4 expression and the corresponding RBPJκ knock out cell line. Results are displayed as in (B). (E). Luciferase reporter assays carried out as in (D) using the RBP-J–dependent C promoter reporter construct.

EBNA2 binds to many target gene enhancers through the cell transcription factor RBPJ (CBF1)(22). We investigated whether EBNA2 activation of *miR-155HG* E2 was mediated via RBPJ. ChIP-QPCR analysis of RBPJ binding in the GM12878 LCL detected RBPJ binding at E2 and not E1, consistent with a role for RBPJ in EBNA2 activation of E2 (Figure 2C). To confirm this, we carried out reporter assays in a DG75 RBPJ knock-out cell line (33). This cell line was derived from a different parental DG75 cell line that also lacks IRF4 expression, so for comparison we also carried out reporter assays in the parental DG75 wild type cell line (Figure 2D). Our data demonstrated that EBNA2 activated transcription of the *miR-155HG* E1 and E2 containing reporter construct in the wild type DG75 cell line to the same extent as the EBV C promoter control, confirming our previous results (Figure 2D). However in DG75 RBPJ knock-out cells, the activation of this reporter construct by EBNA2 was almost completely abolished (Figure 2D). This mirrored the loss of EBNA2 activation observed for the RBPJ-dependent viral C promoter (Figure 2E). These data also provide further evidence that EBNA2 activation of *miR-155HG* E2 is not an indirect effect mediated by IRF4 upregulation and we confirmed that IRF4 expression is not induced by EBNA2 in this cell background (Figure 2D).

Interestingly, EBNA2 binding sites often coincide with binding sites for IRF4 or IRF4-containing transcription complexes, indicating that IRF4 may be involved in EBNA2 binding to DNA (25, 34). However, our results indicate that IRF4 is not required for EBNA2 targeting of *miR-155HG* E2 enhancer element since EBNA2 activation was efficient in the absence of IRF4 (Figure 1D). We conclude that EBNA2 can directly upregulate *miR-155HG* transcription through a distal RBPJ-dependent enhancer (E2) independently of IRF4.

### IRF4 independently activates miR-155HG via promoter and enhancer elements

Our data demonstrate that IRF4 is not required for the effects of EBNA2 on *miR-155HG* transcription. However, IRF4 can independently activate the *miR-155HG* promoter through an ISRE (29). It is not known whether IRF4 can also activate *miR-155HG* transcription through enhancer elements. We therefore tested whether exogenous expression of IRF4 in DG75 cells can activate *miR-155HG* transcription via upstream enhancers. Because IRF4 activates the control plasmid (pRL-TK), Firefly reporter activity was normalized to actin expression as a previously described alternative in these assays (35). Consistent with published data, we found that exogenous expression of IRF4 resulted in a 4-fold increase in *miR-155HG* promoter activity (29). The presence of E1 did not result in any further increase in *miR-155HG* transcription by IRF4 (Figure 3). However, the additional presence of E2 increased the activation of the *miR-155HG* reporter to 10-fold. These data demonstrate that *miR-155HG* E2 is IRF4-responsive and contributes to IRF4 activation of *miR-155HG* transcription.

**FIG 3.**
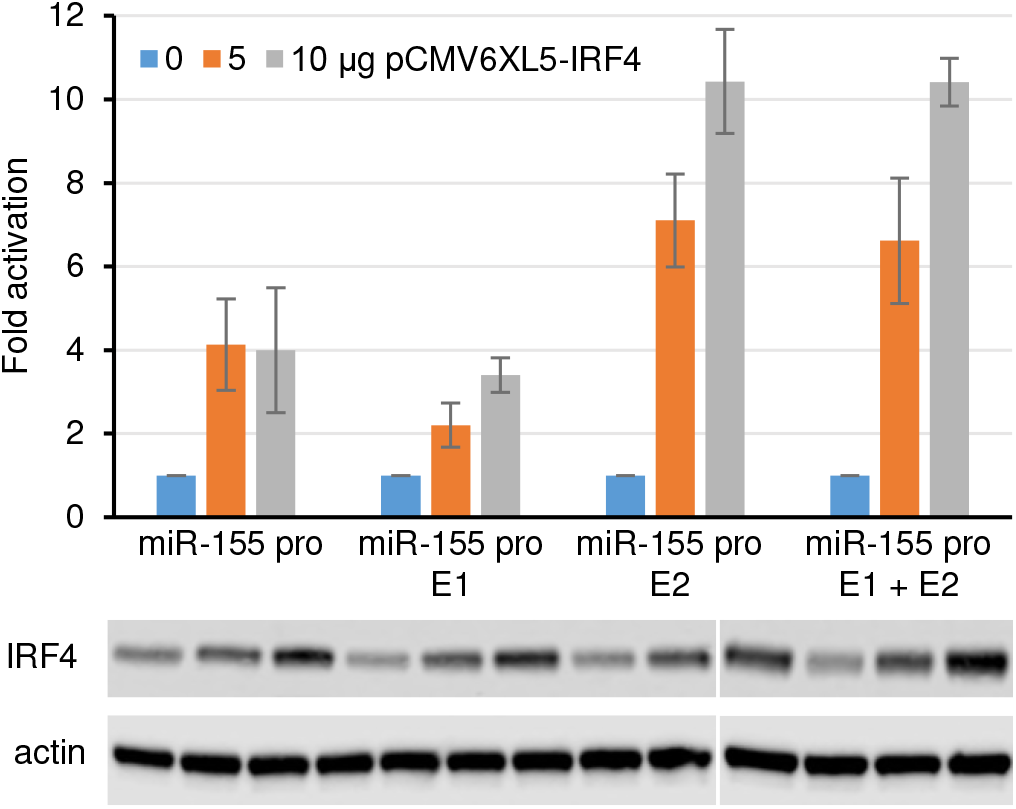
The effects of IRF4 on *miR-155HG* promoter and enhancer elements. DG75 cells were transfected with 2 μg of pGL3 firefly luciferase reporter constructs containing the *miR-155HG* promoter either alone or in the presence of enhancer E1, E2 or both E1 and E2. Assays were carried out in the absence or presence of 5 or 10 μg of the IRF4-expressing plasmid pCMV6XL5-IRF4. Western blot analysis of IRF4 and actin expression is shown below the bar chart. Firefly luciferase signals were normalized to actin western blot signals and fold activation relative to the signal for each construct in the absence of EBNA2 is shown. Results show the mean of three independent experiments plus or minus standard deviation.

Taken together our results indicate that *miR-155HG* promoter activation by IRF4 and the independent effects of IRF4 and EBNA2 on a specific *miR-155HG* enhancer contribute to the high level expression of *miR-155HG* and miR-155 in EBV-infected B cells.

### An IRF4 upstream enhancer is activated by EBNA2 through RBPJ

Our data support a role for IRF4 as a key regulator of *miR-155HG* expression in EBV infected cells. *IRF4* is also an EBNA2 target gene, but the mechanism of *IRF4* upregulation by EBNA2 has not been defined (29, 31). RNA pol II ChiA-PET analysis recently identified a number of upstream regions that interact with *IRF4* in the GM12878 LCL (28). These include the transcription unit of *DUSP22,* an intergenic region upstream from *DUSP22* predicted to be a super-enhancer and intergenic regions between *IRF4* and *DUSP22.* The upstream super-enhancer linked to both *DUSP22* and *IRF4,* so likely represents an important regulatory region (28). EBNA2 ChIP-sequencing data that we obtained using EBV infected cells derived from a Burkitt’s lymphoma cell line additionally identified two large EBNA2 binding peaks within the region 35 kb directly upstream of *IRF4* (Figure 4A). We investigated the potential role of these regions in EBNA2 activation of *IRF4.* These putative proximal and distal EBNA2-bound enhancer regions are referred to as *IRF4* enhancer 1 (E1) and *IRF4* enhancer 2 (E2), respectively (Figure 4A). Luciferase reporter assays carried out in the two different DG75 cell line clones in the absence or presence of transient EBNA2 expression demonstrated that EBNA2 had a small activating effect on the *IRF4* promoter (Figure 4B and D). The presence of *IRF4* E1 reduced basal transcription by 2-fold and increased EBNA2 activation to up to 6.6-fold similar to the level of EBNA2 activation observed for the EBV C promoter (Figure 4B). The additional inclusion of *IRF4* E2 alongside *IRF4* E1 had little further effect on EBNA2 activation (Figure 4B). These data indicate that *IRF4* E1 acts as an EBNA2-responsive enhancer. Consistent with EBNA2 activation through RBPJ, ChIP-QPCR detected RBPJ binding at *IRF4* E1 and not E2 (Figure 4C). Accordingly, EBNA2 activation of the *IRF4* enhancer construct was decreased from 5.7-fold to 1.9 fold in RBPJ knock out cells. Our data therefore demonstrate that EBNA2 can activate *IRF4* transcription through an RBPJ-dependent enhancer (E1) located 13 kb upstream from the transcription start site (TSS).

**FIG 4.**
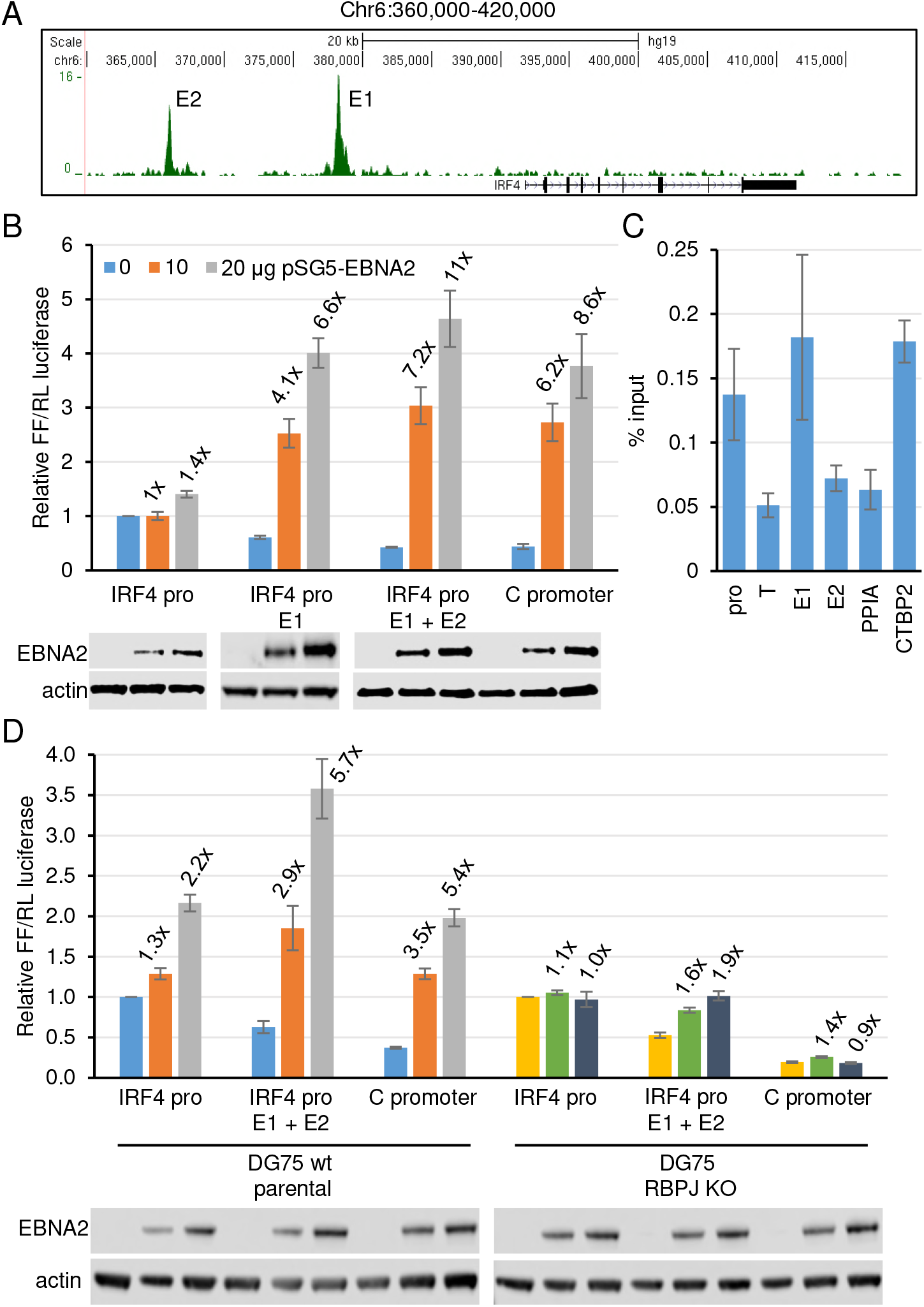
The effects of EBNA2 on *IRF4* promoter and enhancer elements. (**A**) EBNA2 ChIP-sequencing reads at the *IRF4* locus in Mutu III Burkitt’s lymphoma cells (23). The positions of the two main EBNA2-bound putative enhancer regions are indicated (E1 and E2). (**B**) *IRF4* luciferase reporter assays in the presence or absence of EBNA2. DG75 cells were transfected with 2 μg of pGL3 firefly luciferase reporter constructs containing the *IRF4* promoter either alone or in the presence of enhancer E1 or both E1 and E2. Assays were carried out in the absence or presence of 10 or 20 μg of the EBNA2-expressing plasmid pSG5-EBNA2 and 0.5 μg of Renilla luciferase control plasmid (pRL-TK). Firefly luciferase signals were normalized to Renilla luciferase signals and expressed relative to the signal obtained for the *IRF4* promoter in the absence of EBNA2. EBNA2 activation of an EBV C promoter reporter construct was used as a positive control. Results show the mean of three independent experiments plus or minus standard deviation. Fold activation by EBNA2 relative to the signal obtained for each construct in the absence of EBNA2 is shown above each bar. Western blot analysis of EBNA2 is shown below each bar chart, with actin providing a loading control. (**C**) ChIP-QPCR analysis of RBPJκ binding at the *IRF4* locus in GM12878 cells. Precipitated DNA was analysed using primer sets located at the promoter, E1, E2 and in a trough between the promoter and E1 (T). EBNA2 binding at the transcription start site of *PPIA* and at the previously characterised *CTBP2* binding site were used as negative and positive binding controls, respectively. Mean percentage input signals, after subtraction of no antibody controls, are shown plus or minus standard deviation for three independent ChIP experiments. (**D**) Luciferase reporter assays carried out using the *IRF4* promoter, *IRF4* E1 and E2 construct and the RBPJ–dependent C promoter reporter construct in DG75 wt parental cells and the corresponding RBPJκ knock out cell line. Results are displayed as in (B).

### Deletion of miR-155HG E2 from the B cell genome reduces mIR-155HG expression

Since EBNA2 and IRF4 can activate transcription through *miR-155HG* E2 in reporter assays, we next tested the role of this enhancer in the regulation of *mIR-155HG* in EBV infected B cells. To do this, we used CRISPR/Cas9 gene editing to remove the region encompassing E2 (Figure 5A) from the genome of the EBV immortalized LCL IB4. We designed two small guide RNAs (sgRNAs), one targeting a region 5’ to the enhancer and one targeting a region 3 ‘ to the enhancer, so that DNA repair following Cas9 cleavage would generate an E2 deletion (Figure 5A). Both sgRNAs comprised 20 nucleotide sequences that target the genomic region adjacent to a protospacer adjacent motif (PAM) required for Cas9 cleavage (Figure 5C). sgRNAs were transfected into IB4 cells alongside Cas9 protein and single cell clones were generated by limiting dilution. PCR screening was used to identify cell line clones containing E2 deletions using a forward primer located 5’ of the E2 region and a reverse primer located 3’ of E2 to amplify a 180 bp DNA product across the deletion site (Figure 5A and B). This primer set did not amplify DNA from intact templates containing E2 as the amplicon (1.75 kb) was too large for efficient amplification under the conditions used. For three cell line clones tested (C4D, C2B and C5B) we detected amplification of a 180 bp PCR product consistent with the presence of an E2 deletion (Figure 5B). We did not detect this PCR product in parental IB4 cells and an additional clone, C4A indicating that this cell line clone did not contain a deletion (Figure 5B). Sequencing of the PCR products amplified across the deletion site confirmed the E2 deletion (Figure 5C). Clones C4D and C2B contained deletions consistent with cleavage by Cas9 three bases upstream from the PAM sequence as expected, and the subsequent ligation of the cleaved ends. Clone C5D had an additional deletion of 8 nucleotides at the 5’ cut site indicating loss of a small amount of additional DNA during the DNA repair and religation process (Figure 5C).

**FIG 5.**
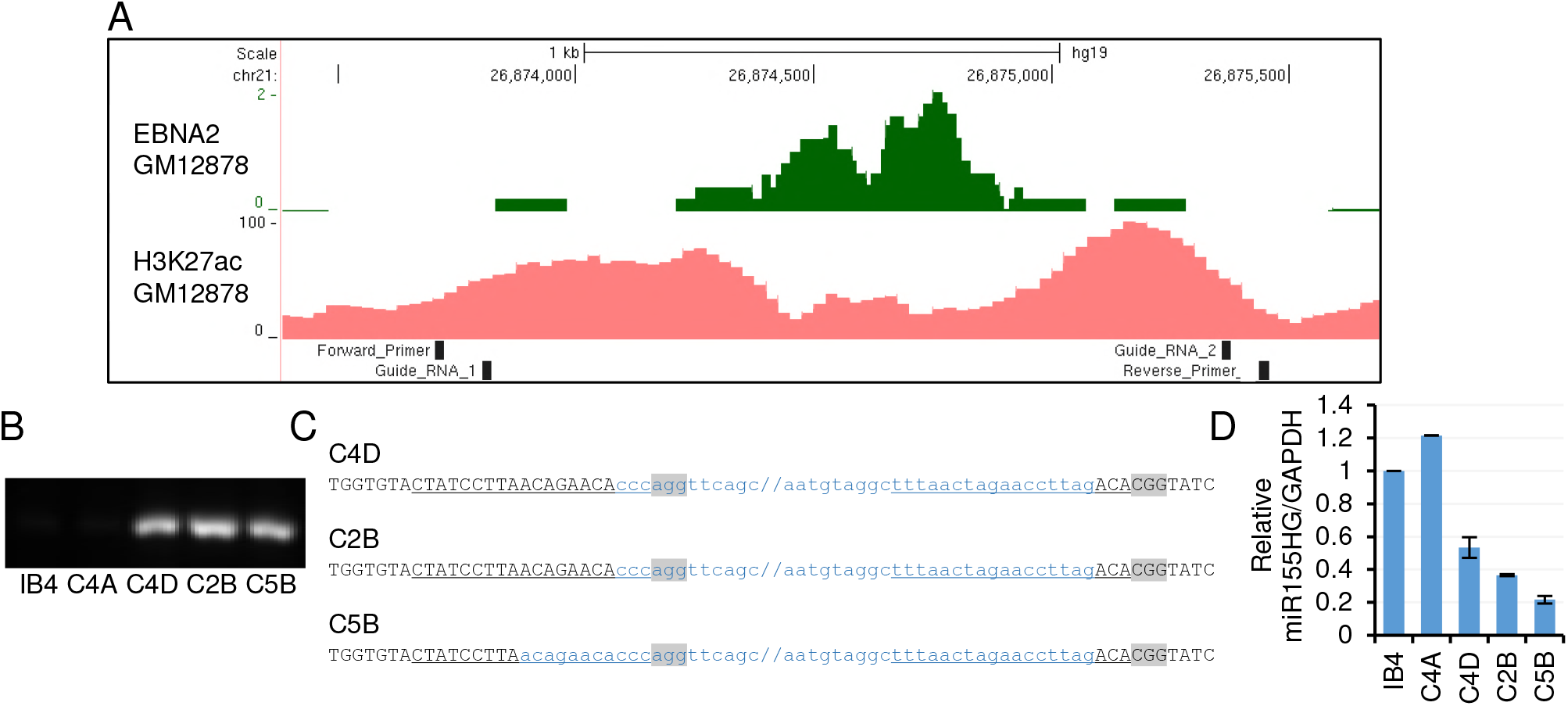
The effects of CRIPSR/Cas9-mediated deletion of *mIR-155HG* enhancer 2. (**A**) EBNA 2 ChIP-sequencing reads in GM12878 cells (24) and H3K27Ac signals in GM12878 from ENCODE at the *miR-155HG* enhancer 2 region. The locations of the guide RNAs used for CRISPR gene editing and the PCR primers used for screening cell clones are indicated. (**B**) PCR analysis of single cell clones obtained by limited deletion following transfection of guide RNAs and Cas 9 protein using primers that span the deletion site and only efficiently amplifies a product (180 bp) from templates carrying an E2 deletion. (**C**) DNA sequence of the deletion spanning PCR products from the C4A, C2B and C5B cell lines. Black uppercase text shows the sequence present in the PCR products and blue lowercase text shows the 5’ and 3’ ends of the deleted region, with forward slashes showing the position of the remaining ~1.5 kb of deleted DNA. The sgRNA target sequences are underlined and PAM sequences are shown in grey. (**D**) QPCR analysis of total RNA extracted from IB4 cells or cell line clones using primers specific for *miR-155HG* and GAPDH. *miR-155HG* signals were normalized by dividing by GAPDH signals and expression levels are shown relative to the signal in IB4 parental cells. Results show the mean -/+ standard deviation of PCR duplicates from a representative experiment.

We next used real-time PCR analysis to determine whether deletion of *miR-155HG* E2 affected the levels of endogenous *miR-155HG* RNA in IB4 cells. We found that all three deletion mutant cell line clones had reduced levels of *miR-155HG* transcripts compared to parental IB4 cells or the non-deleted C4A cell line (Figure 5D). *miR-155HG* RNA expression was reduced by 47%, 63% and 78% in cell line clones C4D, C2B and C5B, respectively (Figure 5D). This indicates that the RBPJ-dependent EBNA2 responsive enhancer (E2) located 60 kb upstream of *miR-155HG* plays an important role in maintaining *miR-155HG* expression in EBV infected cells. Given that miR-155 is derived by processing of the *miR-155HG* transcript, our data indicate that this enhancer would be important in controlling miR-155 expression.

In summary we have identified and characterized new enhancer elements that play a key role in the direct and indirect upregulation of miR-155 expression in EBV infected cells by the EBV transcription factor EBNA2 (Figure 6). Importantly, we show that an EBNA2 and IRF4 responsive enhancer element located 60 kb upstream from the *miR-155HG* TSS is essential to maintain high level *miR-155HG* RNA expression.

**FIG 6.**
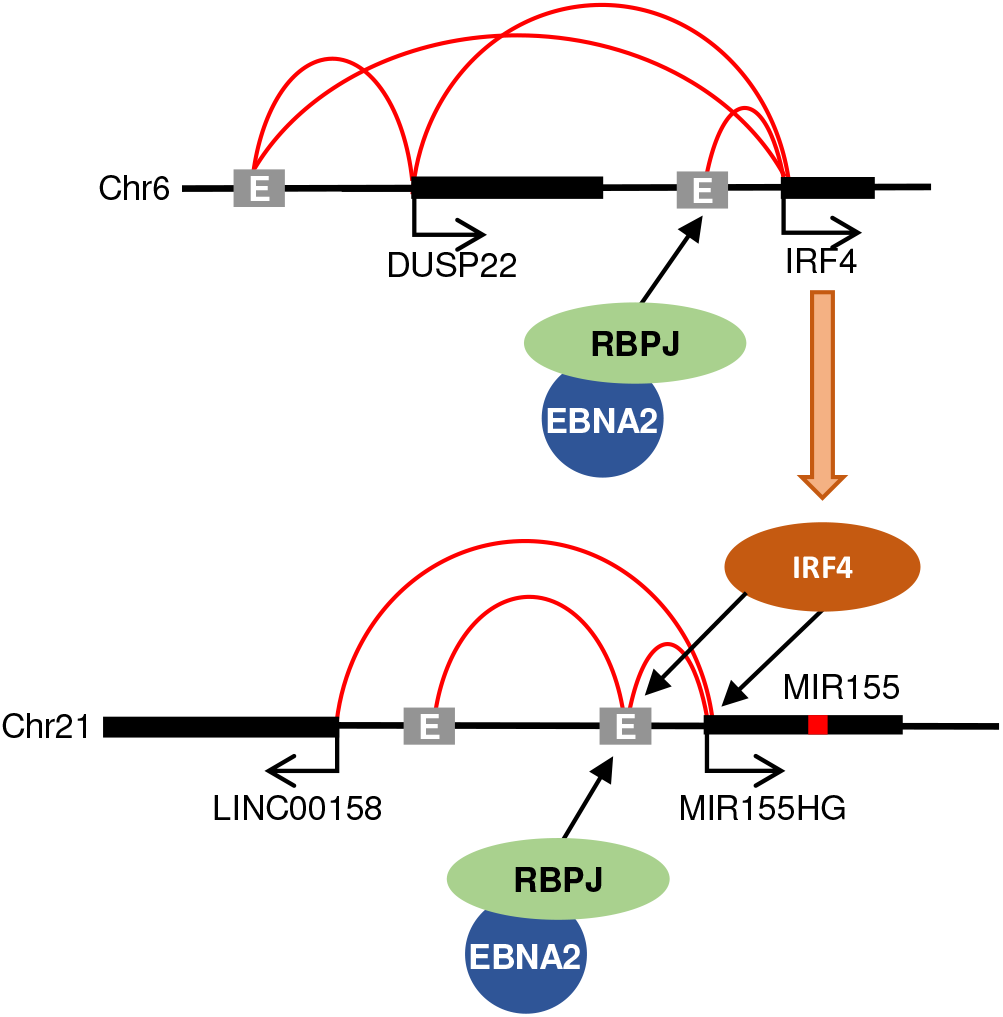
Model for enhancer activation of *IRF4* and *miR155HG* by EBV EBNA2. EBNA2 targets an intergenic enhancer 35 kb upstream of *IRF4* via RBPJ. A super-enhancer upstream of *DUSP22* bound by EBNA 2 also links to both *DUSP22* and *IRF4.* IRF4 then activates *miR-155HG* via the promoter and an intergenic enhancer located 60 kb upstream. EBNA2 activates the *miR-155HG* upstream enhancer via RBPJ. The *miR-155HG* promoter also links to an additional upstream region and the *LINC00158* gene.

## DISCUSSION

We have characterized an enhancer 60 kb upstream of the miR-155-encoding gene *miR-155HG* that is bound by EBNA2, the key transcriptional regulator encoded by Epstein-Barr virus. We have shown that the presence of this enhancer in the B cell genome is required to maintain high level *miR-155HG* expression in an EBV-infected B cell line, indicating that enhancer control is critical for miR-155 upregulation by the virus. This enhancer (enhancer 2) was responsive to EBNA2 in reporter assays and EBNA2 activation was dependent on the expression of host cell protein RBPJ. Since EBNA2 cannot bind DNA directly, this is in line with EBNA2 binding via its interaction with RBPJ (36, 37). *MiR-155HG* enhancer 2 also contains binding sites for a number of other B cell transcription factors (e.g. SPI1 (PU.1), RUNX3, NF-κB rel A, BATF and SRF) that likely play a role in regulating its activity in uninfected B cells. It is also possible that some of these transcription factors may help to stabilize EBNA2 or EBNA2-RBPJ binding in the context of B cell chromatin, a scenario that we cannot examine in reporter assays. PU.1 for example has been shown to bind EBNA2 (38). However, in reporter assays loss of RBPJ alone severely diminishes EBNA2 responsiveness indicating that RBPJ is the major mediator of EBNA2 activation of *miR-155HG* enhancer 2.

MiR-155HG enhancer 2 is located within a region upstream of *miR-155HG* that is detected by CHi-C and RNA pol II ChIA-PET to associate with the *miR-155HG* promoter. Although, another putative enhancer bound by EBNA2 in the GM12878 LCL (enhancer 1) is also present in this region, we found that enhancer 1 was not EBNA2 responsive but did upregulate transcription from the *miR-155HG* promoter in reporter assays. This indicates that this region possesses EBNA2-independent enhancer function. The detected EBNA2 binding at enhancer 1 may therefore be the consequence of looping between enhancer 1 and enhancer 2 that would lead to the precipitation of this region of DNA in EBNA2 ChIP experiments. Interestingly, binding at enhancer 1 is not detected by EBNA2 ChIP-seq in a BL cell background (23), so its activity and looping interactions may be cell-type dependent. Two further upstream regions also interact with the *miR-155HG* promoter by CHi-C and RNA pol II ChIA-PET in LCLs (one intergenic and one proximal to the *LINC00158* promoter). This is consistent with the presence of an active enhancer-promoter hub formed between two intergenic enhancer regions (one of which encompasses enhancer 2) and the promoter-proximal regions of *miR-155HG* and *LINC00158.* In two EBV infected LCL backgrounds (GM12878 and IB4), maximal EBNA2 (and RBPJ) binding at the *miR-155HG* locus is detected in the intergenic interacting region encompassing *miR-155HG* enhancer 2 (22, 24, 28). This is despite the classification of the remaining intergenic region and the *LINC00158* promoter proximal region as EBV super-enhancers based on their chromatin and TF landscape profiles (28). It is therefore possible that EBNA2 accesses the *miR-155HG* enhancer hub and upregulates miR-155 expression through its RBPJ-dependent association with *miR-155HG* enhancer 2. Our observations highlight the importance of testing the EBNA2 responsiveness of EBNA2-bound regions rather than relying on binding profiles alone to assign EBNA2 enhancer function.

The constitutively active EBV membrane protein LMP1 also activates *miR-155HG* transcription. NF-κB and AP-1 sites in the *miR-155HG* promoter have been shown to be important to maintain *miR-155HG* promoter activity in LCLs and two NF-κB sites and the AP-1 site mediate LMP1 responsiveness in transiently transfected EBV negative cells (18, 19). NF-κB RelA also binds to the *miR-155HG* enhancer 2 region and the putative upstream super-enhancer, so it is also possible that LMP1 activation of the NF-κB and AP-1 pathways also activates *miR-155HG* enhancers. Thus promoter (and possibly enhancer) activation by LMP1 and enhancer activation by EBNA2 may all contribute to the high-level miR-155 expression observed in EBV-infected cells.

Our results also revealed that the B cell transcription factor IRF4 can also activate *miR-155HG* transcription via enhancer 2 in addition to its known effects on the *miR-155HG* promoter. IRF4 activates the *miR-155HG* promoter via an ISRE. There are no ISREs within *miR-155HG* enhancer 2, but the 5’ sequence of the PU.1 binding site partially matches a reverse ETS-IRF composite element (EICE), so IRF4 could bind in combination with PU.1. In addition to direct control of miR-155 expression through the *miR-155HG* enhancer, EBNA2 also indirectly influences miR-155 expression through the transcriptional upregulation of *IRF4.* We demonstrate that again enhancer control by EBNA2 plays an important role in *IRF4* activation. In addition to the presence of an EBNA2-bound superenhancer upstream of the neighboring *DUSP22* gene (28), we found that EBNA2 can also upregulate *IRF4* transcription through an RBPJ dependent enhancer located in an intergenic region 35 kb upstream from IRF4. At *IRF4* and *DUSP22,* EBNA2 therefore likely targets multiple enhancers and super-enhancers.

MiR-155 is overexpressed in many tumor contexts, including hematological malignancies and is implicated in cancer therapy resistance (11). It therefore represents an important therapeutic target. The first in human phase I trial of a synthetic locked nucleic acid anti-miR to miR-155 has been initiated and preliminary results show that the inhibitor is well tolerated in patients with cutaneous T cell lymphoma when injected intratumorally (39). Inhibition of miR-155 expression through indirect transcriptional repression has also been tested in acute myeloid leukemia cells using an inhibitor of the NEDD8-activating enzyme (40). NEDD8-dependent ubiquitin ligases regulate NF-κB activity and their inhibition by MLN4924 in AML cells results in reduced binding of NF-κB to the *miR-155HG* promoter and a reduction in miR-155 expression. In mice engrafted with leukemic cells, MLN4924 treatment reduced miR-155 expression and increased survival. These data provide evidence for transcriptional inhibition of miR-155 as a therapeutically viable strategy.

The sensitivity of super-enhancers to transcriptional inhibitors is also being exploited as a therapeutic strategy in various tumor contexts. Super-enhancers often drive the high-level expression of oncogenes and super-enhancer inhibition by CDK7 and BET inhibitors can effectively block tumor cell proliferation and enhance survival in mouse models of disease (41-43). MiR-155 expression in human umbilical vein endothelial cells is sensitive to inhibition by BET and NF-κB inhibitors (44). This was proposed to result from inhibition of an upstream miR-155 super-enhancer, but the region examined actually represents the *miR-155HG* transcription unit, which has high-level histone H3 K27 acetylation (used as a super-enhancer marker) throughout its length when *miR-155HG* is transcriptionally active.

Nonetheless, the study highlights the usefulness of transcription inhibitors in reducing miR-155 expression. Our identification and characterisation of the enhancers that drive *miR-155HG* transcription in B cells may therefore open up new therapeutic opportunities for the inhibition of miR-155 expression in numerous B cell cancer contexts where miR-155 is a key driver of tumor cell growth.

## METHODS

### Cell lines

All cell lines were cultured in RPMI 1640 media (Invitrogen) supplemented with 10% Fetal Bovine serum (Gibco), 1 U/ml penicillin G, 1 μg/ml streptomycin sulphate and 292 μg/ml L-glutamine at 37°C in 5% CO2. Cells were routinely passaged twice-weekly. The DG75 cell line originates from an EBV negative BL (45). DG75 cells cultured in our laboratory (originally provided by Prof M. Rowe) express low levels of IRF4, but DG75 cells obtained from Prof B. Kempkes (referred to here as DG75 wt parental cells) lack IRF4 expression. The DG75 RBPJ (CBF1) knock-out cell-line (SM224.9) was derived from DG75 wt parental cells (33). (46). IB4 (47) and GM12878 (obtained from Coriell Cell Repositories) are EBV immortalised lymphoblastoid cell lines (LCLs) generated by infection of resting B cells *in vitro*

### Plasmid construction

The *miR-155HG* promoter sequence from -616 to +515 (Human GRCh37/hg19 chr 21 26933842-26934972) was synthesized by GeneArt Strings^®^ (Invitrogen) to include XhoI and HindIII restriction enzyme sites and cloned into pGL3 basic (Promega) to generate the pGL3 miR155HG promoter construct. The pGL3miR-155HG enhancer 1 (E1) construct was generated in a similar way by synthesis of the promoter and upstream E1 region (chr21 26884583-26885197) as a single DNA fragment that was then cloned into pGL3 basic. To generate the miR-155HG promoter E1 + E2 construct, the promoter and E1 and E2 regions (chr21 26873921-26875152) were synthesized as a single DNA fragment and cloned into pGL3 basic. The pGL3-miR-155HG promoter E2 construct was generated using sequence and ligation independent cloning. The E2 region was amplified by PCR from the miR-155HG promoter E1 + E2 construct using primers containing vector and insert sequences (forward 5’ TCTTACGCGTGCTAGCCCGGGCTCGAGGAGAGGTTTAAAGCACTCAGACAGC 3’ and reverse 5’ GGGCTTTGAGAACGTTTGTACCTCGAGGATCTAGAACCTCTGGAGTTGGAGA T 3’). The pGL3-miR-155HG promoter vector was digested with XhoI and then T4 DNA polymerase was used to further resect the cut ends to allow the insert to anneal to extended single-stranded regions of the vector. Single-strand DNA gap filling occurred through DNA repair following transformation of the plasmid into *E.coli.*

The *IRF4* promoter sequence from -739 to +359 (Human GRCh37/hg19 chr6 391024392121) was synthesized by GeneArt (Invitrogen) and the promoter fragment was amplified from the supplied vector (pMK-RQ) using primers to introduce XhoI restriction sites at each end (forward 5’ GTCTCGAGATTACAGGCTTGAGCCACA 3’, reverse 5’GACTCGAGCTGGACTCGGAGCTGAGG 3’). The promoter was then cloned into the XhoI site of pGL3 basic (Promega) to generate the pGL3 IRF4 promoter construct. *IRF4* enhancer 1 (E1) (chr6 377854-379089) was amplified from genomic DNA using primers to introduce NheI and XhoI sites (5’ forward GAGCTAGCATCGCTTGAGGTTGCAGTG 3’ and reverse 5’ GTCTCGAGTGAAGCAGGCACTGTGATTC 3’). The XhoI site was end filled using Klenow and the E1 fragment was cloned upstream of the promoter into the NheI and SmaI sites of the pGL3 IRF4 promoter construct. E2 (chr6 365659-366654) was amplified by PCR using primers designed to introduce SacI and NheI sites (forward 5’ GAGAGCTCAGCCATCTCCATCATCTGGT 3’ reverse 5’ GAGCTAGCATGTGGAACGCTGGTCC 5’) and cloned upstream of E1 into the SacI and NheI sites of the pGL3 IRF4 promoter E1 construct.

### Luciferase reporter assays

DG75 cell lines were electroporated with plasmid DNA at 260 V and 950 μF (BioRad Gene Pulser II) using 0.4cm cuvettes and luciferase assays carried out as described previously with some modifications (48). Briefly, DG75 cells were diluted 1:2 into fresh medium 24 hours prior to electroporation. For transfection, cells were pelleted and conditioned media reserved for later use. Cells were then resuspended in serum-free media to a density of 2×10^7^ cells/ml. 500 μl of cell suspension was pre-mixed with DNA and then added to the cuvette and immediately electroporated. Transfected cells were then transferred to 10 ml of pre-warmed conditioned media, and cultured for 48 hours in a humidified incubator at 37°C, with 5% CO2.

Cells were transfected with 2μg of the pGL3 luciferase reporter plasmids and 0.5μg pRL-TK (Promega) as a transfection control where indicated. Transfection reactions also included 10 or 20 μg of the EBNA2 expressing plasmid (pSG5 EBNA2), 5 or 10 μg of IRF4 expressing plasmid (pCMV6XL5-IRF4, Cambridge Biosciences) or empty vector control. One tenth of each transfection was processed for Western blotting to analyse EBNA2, IRF4 and actin protein expression levels. The remaining cells were lysed and firefly and Renilla luciferase activity measured using the dual luciferase assay (Promega) and a Glowmax multi detection system (Promega). For transfections where IRF4 was expressed, firefly luciferase signals were normalized to actin expression.

### CRISPR

CRISPR guides were designed using www.benchling.com_to excise *miR-155HG* enhancer 2 from the B cell genome in the IB4 LCL by targeting genomic regions located 5’ and 3’ of the enhancer. Guides were selected that had an on-target and off-target score that was above 60% (26873822 CTATCCTTAACAGAACACCC and 26875376 TTTAACTAGAACCTTAGACA) and then ordered as TrueGuide Modified Synthetic sgRNAs from GeneArt (Invitrogen). IB4 cells were diluted 1:2 into fresh medium 24 hours prior to transfection. 1×10^6^ cells were then washed in PBS, pelleted and resuspended in 25 μl of resuspension Buffer R (Invitrogen). The guide RNA and Cas9 mix was prepared by adding 7.5 pmol of GeneArt TrueCut Cas9 Protein V2 (Invitrogen) and 7.5 pmol of sgRNAs to 5 μl of resuspension Buffer R and incubating at room temperature for 10 minutes. 5 μl of cell suspension (2 x10^5^ cells) was then mixed with 7ul of the Cas9/sgRNA complex. 10ul of the cell Cas9/sgRNA mix was then electroporated using the Neon transfection system (Invitrogen) at 1700V for 20 ms with 1 pulse. Transfections were carried out in duplicate and electroporated cells were immediately transferred to two separate wells of a 24 well plate containing 0.5 ml of pre-warmed growth media. Cells were kept in a humidified incubator at 37°C, with 5% CO2 for 72 hours. Cells were then sequentially diluted over a period of 2 weeks and subject to limited dilution in 96 well plates to obtain single cell clones. Cell line clones were screened by PCR for genomic deletion using the PHIRE Tissue Direct PCR Master Mix kit (Thermo Scientific). The forward primer (5’ AAATTCCGTGGCTAGCTCCA 3’) hybridized to a region 5’ of enhancer 2 and two different reverse primers targeted either a region within enhancer 2 (reverse primer 1, 5’ AATGGGATGGCTGTCTGAGT 3’) or a region 3’ to enhancer 2 (reverse primer 2, 5’ CTGCTAAGGGAATGTTGAACAAA 3’). Deletions were confirmed by DNA sequencing of the PCR product generated using the forward PCR primer.

### SDS-PAGE and Western Blotting

SDS-PAGE and Western blotting was carried out as described previously (48, 49) using the anti-EBNA2 monoclonal antibody PE2 (gift from Prof M. Rowe) anti-actin 1/5000 (A-2066, Sigma) and IRF4 1/2000 (sc6059, Santa Cruz). Western blot visualization and signal quantification was carried out using a Li-COR Imager.

### ChIP-QPCR

ChIP-QPCR for RBPJ was carried out as described previously (24). *MiR-155HG* locus primers were located in the *miR-155HG* promoter (forward 5’ AGCTGTAGGTTCCAAGAACAGG 3’ and reverse 5’ GACTCATAACCGACCAGGCG 3’, *miR-155HG* enhancer 1 (forward 5’ ACCTGTTGACTTGCCTAGAGAC 3’ and reverse 5’ TTCTGGTCTGTCTTCGCCAT 3’), a ‘trough’ region between *miR-155HG* enhancer 1 and enhancer 2 (forward 5’ TATTCAGCTATTCCAGGAGGCAG 3’ and reverse 5’ GTGACATTATCTGCACAGCGAG 3’), and *miR-155HG* enhancer 2 (forward 5’ CCTAGTCTCTCTTCTCCATGAGC 3’ and reverse 5’ AGTTGATTCCTGTGGACCATGA 3’). *IRF4* locus primers were located in the *IRF4* promoter (forward 5’ TCCGTTACACGCTCTGCAA 3’ and reverse 5’ CCTCAGGAGGCCAGTCAATC 3’), a ‘trough’ region between the *IRF4* promoter and enhancer 1 (forward 5’ TGTGACAAGTGACGGTATGCT 3’ and reverse 5’ TTGTAACAGCGCCTAATGTTGG 3’), *IRF4* enhancer 1 (forward 5’ TTACCACCTGGGTACCTGTCT 3’ and reverse 5’ ACAGTAGCATGCAGCACTCTC 3’) and *IRF4* enhancer 2 (forward 5’ AGTGAGACGTGTGTCAGAGG 3’ and reverse 5’ AAGCAGGCACTGTGATTCCA 3’).

### RT-QPCR

Total RNA was extracted using TriReagent (Sigma) and RNA samples then purified using the RNeasy kit (Qiagen). RNA concentrations were determined using a Nanodrop 2000 (Thermo Scientific) and 1 μg was used to prepare cDNA using the ImProm II reverse transcription kit with random primers (Promega). Quantitative PCR was performed in duplicate using the standard curve absolute quantification method on an Applied Biosystems 7500 real-time PCR machine as described previously (23) using published QPCR primers for *mIR-155HG* (BIC) (29) (forward 5’ACCAGAGACCTTACCTGTCACCTT3’ and reverse 5’GGCATAAAGAATTTAAACCACAGATTT 3’) and GAPDH (forward 5’ TCAAGATCATCAGCAATGCC 3’ and reverse 5’ CATGAGTCCTTCCACGATACC 3’)

### Capture Hi-C

Previously described capture Hi-C data from GM12878 and CD34+ cells were examined for interactions that were captured using baits comprising a 13,140 bp HindIII fragment encompassing the *mIR-155HG* promoter (GRCh38/hg19 chr21:26926437-26939577) and a 2,478 bp HindIII fragment that encompasses the miR-155 genomic sequence in exon 3 (GRCh38/hg19 chr21:26945874-26948352).

## ACKNOWLEDGEMENTS

This work was funded by the Medical Research Council (MR/K01952X/1) to MJW and Bloodwise (15024 to MJW and 14007 to CSO). Funding for open access charge: Medical Research Council.

